# Hepatotoxicity assessment of antimalarial compounds from microorganisms using a liver organ-on-a-chip system

**DOI:** 10.64898/2026.09.10.750749

**Authors:** Nadeeya Mad-adam, Htun Aung Kyaw, Intira Poonsawaeng, Pimonrat Ketsawatsomkron, Nantiya Bunbamrung, Pattama Pittayakhajonwut, Chitti Thawai, Kenjiro Muta

## Abstract

Malaria remains a critical public health threat in sub-Saharan Africa and Southeast Asia owing to the emergence of resistance to gold-standard artemisinin-based combination therapies. This epidemiological shift drives an urgent need to explore novel bioactive compounds for malaria treatment. Microorganisms produce diverse secondary metabolites with potent biological activities. To develop microbe-derived antimalarial drugs, rigorous safety evaluations are crucial, particularly regarding hepatotoxicity. Therefore, we evaluated the hepatotoxic potential of three compounds with antimalarial attributes: thiolutin (NATPP0650), cochliodinol (NATPP0437), and 4’-hydroxy-mycophenolic acid (NATPP0604). Acute hepatotoxic responses were determined by biochemical assays for cell viability and damage in HepG2 cells following 24 h exposure, comparing conventional two-dimensional cultures with an organ-on-a-chip (OOC) system featuring in vivo-like functionality and long-term culture capabilities. In both culture platforms, NATPP0650 and NATPP0437 caused significant toxicity at concentrations of 10 and 50 µM, whereas NATPP0604 was relatively non-toxic. Based on the OOC data, NATPP0650 and NATPP0437 exhibited CC_50_ values of 3.04 and 2.21 µM, respectively, whereas NATPP0604 had a CC_50_ value > 50 µM, offering a superior safety profile with a selectivity index > 23.7. Functional assessment of HepG2 OOCs showed that NATPP0650 and NATPP0437 markedly suppressed albumin production at 10 and 50 µM, whereas NATPP0604 showed minimal inhibition. However, long-term (7 days) toxicity testing on the OOC model revealed a critical finding: NATPP0604 exerted delayed hepatotoxic effects that were undetectable in the 24-h study, as evidenced by partial reductions in cell viability at 10 and 50 µM and decreased albumin production across all tested concentrations. These findings demonstrate the therapeutic safety potential of NATPP0604 for further development as an antimalarial drug and underscore the importance of the long-term OOC data in establishing accurate safety margins for future clinical trials.

## Introduction

Malaria is a mosquito-borne disease caused by *Plasmodium* parasites. In 2024, the estimated number of malaria cases across 80 endemic countries was 282 million, representing an increase of approximately 9 million cases from 2023 [1]. Moreover, global malaria deaths rose by approximately 12,000 from 2023 to 2024, resulting in an estimated 610,000 deaths in 2024 [1]. Critically, this high burden persists despite the widespread deployment of effective prevention tools, such as insecticide-treated nets, and first-line treatments such as artemisinin (ART)-based combination therapies (ACTs), respectively [1].

Malaria infection begins when *Plasmodium* sporozoites are transmitted to the human host through the bite of an infected female *Anopheles* mosquito [2, 3]. Upon entering the bloodstream, sporozoites rapidly migrate and infect hepatocytes [2, 3]. This initiates the asymptomatic liver stage, during which the parasites undergo extensive asexual replication and differentiation into schizonts, ultimately producing thousands of merozoites [2, 3]. Approximately 6 to 16 days after the initial infection, the infected hepatocytes rupture, releasing these merozoites into the systemic circulation [2, 3]. The released merozoites then invade red blood cells, commencing the symptomatic blood-stage infection cycle [2, 3]. The rapid replication and sustained destruction of red blood cells during this stage lead to the clinical manifestations of malaria, such as severe anemia, respiratory distress, and cerebral malaria, all of which require immediate medical intervention [4, 5].

ACTs are currently used as first-line treatments for malaria in endemic countries worldwide [6]. The World Health Organization recommends six ACT regimens for treating malaria: artemether-lumefantrine, artesunate-amodiaquine, artesunate-mefloquine, dihydroartemisinin-piperaquine, artesunate-sulfadoxine-pyrimethamine, and artesunate-pyronaridine [1]. However, the efficacy of these cornerstone treatments is increasingly threatened by the emergence of drug resistance [1, 7–9]. Partial resistance to ART has been frequently detected in both African and Southeast Asian countries, posing a significant global challenge to malaria control [1, 7–9]. Furthermore, ART itself is known to cause various adverse effects, such as post-ART (or post-artesunate) delayed hemolysis, with incidences as high as 22% in malaria patients, and dose-dependent prolongation of the QT interval [10, 11]. Moreover, repeated treatments with artemether-lumefantrine or artesunate-amodiaquine have been shown to reduce the viability of HepG2, a commonly used liver cell line, thereby limiting the use of ACTs in prophylaxis [12]. Therefore, there is an urgent need to develop alternative, safe, and effective antimalarial drugs.

Natural products remain crucial sources of biologically active scaffolds for drug discovery. Isolated from a wide range of organisms spanning from bacteria to higher eukaryotes, these compounds have been used in human disease management for centuries [13, 14]. According to Bérdy [15], approximately 40% of known microbial biological compounds are derived from fungi, and approximately 45% originate from actinomycetes. From our depository of microbe-derived bioactive compounds, we selected three antimalarial agents to investigate their potential hepatotoxicity: thiolutin (NATPP0650) isolated from *Streptomyces marinisediminis* JHD1^T^ [16], cochliodinol (NATPP0437) isolated from *Chaetomium globosum* [17], and 4′-hydroxy-mycophenolic acid (4’-hydroxy-MPA, NATPP0604) isolated from *Penicillium parvum* BCC75476 (unpublished data); all three possess documented antimalarial activity, with the half-maximal inhibitory concentration (IC_50_) values of 5.92, 4.39, and 2.11 µM (equivalent to 1.35, 2.22, and 0.71 µg/mL), respectively [16, 17]. These values fall within the established threshold for active antimalarial hits (IC_50_ of 1−20 µM for good activity [18]. Additionally, NATPP0650, NATPP0437, and NATPP0604 showed cytotoxicity against African green monkey kidney epithelial (Vero) cells [19], with 50% cytotoxic concentration (CC_50_) values of 2.86, 2.13, and 33.9 µM, respectively [16, 17]. Based on these preliminary data, NATPP0604 appears to be a promising antimalarial candidate owing to its favorable efficacy-to-toxicity ratio.

Drug development requires a substantial investment of time and resources [20]. Nevertheless, clinical trial failures and post-approval withdrawals continue to inflict significant setbacks on the advancement of new therapies [21]. Notably, 22% of clinical trial failures and 32% of market withdrawals have been attributed to drug-induced hepatotoxicity [20]. In antimalarial drug development, adverse effects commonly associated with elevated liver enzymes are the primary bottleneck limiting candidates from progressing through the discovery pipeline [22, 23]. While animal models have been instrumental in malaria research, they exhibit notable physiological differences from the human pathology of *Plasmodium* parasites, such as the life cycle and chronicity of infection, limiting their translational relevance [24]. This discrepancy emphasizes the need to apply more human-relevant preclinical models in the drug development process [25]. Recently, under the Food and Drug Administration Modernization Act 3.0, alternatives to animal testing, including advanced in vitro models and computational simulations, have been approved for safety and efficacy assessments of novel drugs and biological products [26].

In toxicity testing, conventional two-dimensional (2D) cultures have been used for decades owing to their cost-effectiveness and simple methodology [27]. However, the static nature of 2D cultures and their limited cell–cell and cell–extracellular matrix (ECM) interactions make them less representative of in vivo environments [28, 29]. Moreover, rapid cell proliferation and subsequent contact inhibition-induced senescence upon reaching confluence restrict the utility of 2D cultures for long-term toxicity assessments [30]. Note that current antimalarial therapies require extended treatment periods, typically 3–7 days or more depending on the parasite species [1, 31–34]. These limitations can be overcome by leveraging organ-on-a-chip (OOC) systems, which incorporate a three-dimensional scaffold, an ECM compartment, and fluidic microchannels [30, 35–37]. These features enable OOCs to recapitulate in vivo-like tissue architecture and function [30] while supporting long-term culture stability [36]. In in vitro liver models, primary human hepatocytes are regarded as the gold standard for predicting drug-induced hepatotoxicity [38]. Nevertheless, given their drawbacks related to inconsistent sample extraction, donor-to-donor variability, and de-differentiation, human hepatic cell lines (such as HepG2 and HepaRG) remain the most widely applied models for malaria replication and drug toxicity screening [30, 36, 39, 40]. HepG2 cells cultured in MIMETAS OrganoPlate OOCs (HepG2 OOCs) have been shown to sustain high viability for up to 14 days [41]. Furthermore, HepG2 OOCs maintain essential hepatic functions, such as albumin and urea synthesis, significantly better than 2D platforms for at least 14 days [41, 42]. Thus, metabolic and functional readouts can be monitored over extended periods in this OOC model to accurately identify drug-induced hepatotoxic risks [41, 42]. The suitability of HepG2 OOCs for long-term toxicity screening was previously validated by our group during a 7-day hepatotoxic assessment of plant extracts [43]. Relative to sophisticated but convoluted liver OOC models that require unidirectional medium flow and multiple non-parenchymal cell types [44–46], HepG2 OOCs have lower technical complexity, offering a practical, scalable, and high-throughput screening solution for studying drug-induced hepatotoxicity in a more biomimetic and reliable setting.

Here, the HepG2 OOC platform was used to evaluate the hepatotoxic profiles of three antimalarial compounds. Some tests were replicated in conventional 2D cultures for comparison. We hypothesized that the HepG2 OOC system would provide sensitive detection of the hepatotoxicity induced by these compounds. To test this hypothesis, we exposed OOC and 2D cultures of HepG2 cells to the compounds for 24 h and measured the cell viability and cellular damage. Moreover, we evaluated the effect of these three compounds on hepatic functionality by quantifying albumin and urea production within the OOCs. Taking advantage of the extended culture capabilities of the HepG2 OOC platform, we also examined the long-term cytotoxicity and functional disturbances triggered by these compounds over 7 days.

## Materials and Methods

### Source and preparation of the test compounds

NATPP0650, NATPP0437, and NATPP0604 were obtained from distinct microbial sources. NATPP0650 was obtained from the crude whole-cell extract of *Streptomyces* sp. JHD1^T^ via several chromatographic techniques, as previously described by Pansomsuay et al. [16]. NATPP0437 was obtained from the crude extract of the endophytic fungus *C. globosum* BCC 71876 via a Sephadex LH20 column (Cytiva Sweden AB, USA) and preparative high-performance liquid chromatography (HPLC, DIONEX ultimate 3000), as described by Bunbamrung et al. [17]. NATPP0604 was isolated from the crude extract of the fungus *P. parvum* BCC 75476 (Unpublished data) via preparative HPLC. The chemical structures of all three compounds were identified by standard spectroscopic techniques, including NMR spectroscopy, mass spectrometry, and UV (JASCO V730) and IR (ALPHA FT-IR, Bruker) spectrophotometry [16, 17, 47]. The chemical structure of NATPP0604 was confirmed by spectroscopic comparison with previously reported 4’-hydroxy-MPA [47]. HRESIMS spectra were recorded using a MicrOTOF mass spectrometer (Bruker) to determine the precise molecular mass. NMR spectra were obtained using either a Bruker Avance III HD 400 spectrometer or a Bruker Avance III HD 500 MHz spectrometer, and chemical shifts were reported in ppm relative to the internal deuterated solvent standard. HPLC-based purifications were performed using a DIONEX Ultimate 3000 system (Thermo Fisher Scientific, USA) equipped with either Capcell Pak C18 MGII (Shiseido, Japan) or SunFire C18 (Waters, USA) columns, as previously described [16, 17]. A linear gradient system of water and MeCN (Fisher Scientific Korea Ltd., Korea) served as the mobile phase, and UV detection was set at 210 nm. The purity of each compound was confirmed to be > 99% by analytical HPLC.

### Chemical preparation

Troglitazone (Enzo Life Sciences, USA) and the antimalarial compounds (NATPP0650, NATPP0437, and NATPP0604) were dissolved in 100% anhydrous dimethyl sulfoxide (DMSO; Sigma-Aldrich, USA) to prepare 200 and 50 mM stock solutions, respectively. The stocks were aliquoted and stored at −20 °C for one-time use, to prevent freeze-thaw cycles.

### Preparation of the 2D HepG2 cell cultures

HepG2 human hepatocarcinoma cells (HB-8065) were purchased from the American Type Culture Collection (USA). The cells were grown in 100 mm culture dishes with the complete medium comprising Dulbecco’s Modified Eagle’s Medium (Gibco, USA), 10% fetal bovine serum (Sigma-Aldrich, USA), 1 mM sodium pyruvate (Gibco, USA), and 2 mM GlutaMAX (Gibco, USA), using a humidified 5% CO_2_ incubator at 37 °C. For toxicity testing, cells were seeded into a 96-well plate at 25,000 cells/well in the complete medium. After 48 h of incubation, the medium was replaced with William’s E (WE) medium (Gibco, US) supplemented with 10% fetal bovine serum and 2 mM L-glutamine (Gibco, US). After 24 h of acclimation, 2D HepG2 cells were exposed to 0.1% DMSO (vehicle), incremental concentrations (0.001, 0.01, 0.1, 1, 10, and 50 µM) of each compound, or 200 μM troglitazone (positive control) for 24 h. The conditioned medium was collected and stored at −80 °C for subsequent analysis.

### Preparation of the HepG2 OOCs

HepG2 OOCs were prepared according to our previously established protocol [42]. Briefly, HepG2 cells were suspended in Cultrex Collagen I (4 mg/mL, rat tail; Bio-Techne, USA) at 1 × 10^7^ cells/mL. The suspension was infused into the gel inlet chamber of a 2-lane OrganoPlate (cat# 9605-400-B, MIMETAS, The Netherlands). According to the dimensions of the chamber, the seeded number of HepG2 cells in the ECM gel adjacent to the medium path was estimated to be approximately 5,000 cells. Complete WE medium (100 μL) was added to the medium chambers and replaced daily. The OrganoPlate was incubated on an OOC rocker (MIMETAS, The Netherlands) that slanted side-to-side at 7° every 8 min in a humidified 5% CO_2_ incubator at 37 °C. After 3 days of incubation, the HepG2 OOCs were exposed to the test compounds under the same treatment conditions used for the 2D cultures, for either a 24-h or a 7-day exposure window.

### WST-8 assay

Cell viability was assessed using the WST-8 assay [48]. For HepG2 OOCs, the medium chambers were washed five times with serum-free WE medium to remove any residual test compounds that might interfere with the WST-8 assay reaction [49]. The OOC cells were then incubated with 50 μL of WST-8 solution (Abcam, UK) for 3 h on the rocker at 37 °C in the CO_2_ incubator. For the 2D HepG2 cultures, each well was washed twice with serum-free WE and incubated with 50 μL of WST-8 solution for 30 min at 37 °C in the CO_2_ incubator. The absorbance of the metabolized WST-8 solution from both cultures was measured at 450 nm using a microplate reader (BioTek Cytation 5, Agilent Technologies, Inc., USA).

### Lactic dehydrogenase (LDH) assay

To assess cellular damage caused by the test compounds in HepG2 cells, the enzymatic activity of lactate dehydrogenase (LDH) in the conditioned medium was evaluated using an LDH Activity Assay Kit (Sigma-Aldrich, USA) [50]. Following the manufacturer’s protocol, LDH activity was calculated from optical density measurements at 450 nm using a microplate reader every 5 min until the absorbance value of the most active sample exceeded that of the standard sample at the highest concentration.

### Albumin and urea assays

To investigate the effect of the test compounds on hepatic functions, the amounts of albumin and urea in the culture medium were quantified using a Human Albumin ELISA Kit (Abcam, USA) and a Urea Assay Kit (Sigma-Aldrich, USA), respectively. The assays were performed according to the manufacturers’ instructions.

### Selectivity index (SI) calculation

The selectivity index (SI) was calculated to determine the safety of a compound toward HepG2 cells relative to its efficacy against *Plasmodium falciparum* [51–53]. The SI value is defined as the ratio of the CC_50_ against HepG2 cells to the IC_50_ against the *P. falciparum* K1 strain. Each SI value was calculated using the formula: SI = CC_50_/IC_50_. According to the previous reports [51–53], a compound is considered selective toward *Plasmodium falciparum* when the SI is more than 10.

### Robust Z’-factor and power analysis calculations

Robust Z’-factors were calculated using a previously described method [54, 55]. An experiment with a Robust Z’-factor > 0.2 is considered good, whereas a Robust Z’-factor > 0.5 is considered excellent for high-throughput screening. A post hoc power analysis was performed using G*Power version 3.1.9.7 (Heinrich Heine University, Düsseldorf, Germany). The analysis was based on a two-tailed independent samples *t*-test using the observed effect size (Cohen’s d), an α level of 0.05, and the actual sample sizes in each group [56]. Statistical power values > 0.80 (80%) were considered to indicate adequate power for detecting treatment-related differences [57].

### Statistical analysis

All data were produced by performing at least three independent experiments (biological replicates), each consisting of a single well or multiple wells per treatment condition (technical replicates). Data are presented as the mean ± standard error of the mean (SEM). Statistical analyses were performed using GraphPad Prism software version 11 (GraphPad, USA). One-way analysis of variance (ANOVA) followed by Tukey’s post hoc test was used to determine pairwise significance. A p-value < 0.05 was considered statistically significant.

## Results

### Characterization of the 2D and OOC HepG2 cultures for toxicity testing

Prior to evaluating the hepatotoxicity of the three test compounds, we characterized the toxicological responses of HepG2 cells cultured in 2D and OOC platforms using troglitazone, a known human hepatotoxin [58]. This step validated the capacity of each culture platform to detect an established human hepatotoxic agent. The morphological and biochemical responses of 2D and OOC HepG2 cultures were determined following exposure to 200 µM troglitazone for 24 h. In both platforms, HepG2 cells became smaller and more rounded than those of the vehicle controls (Fig 1A), implying a distinct toxic effect.

**Fig 1.**
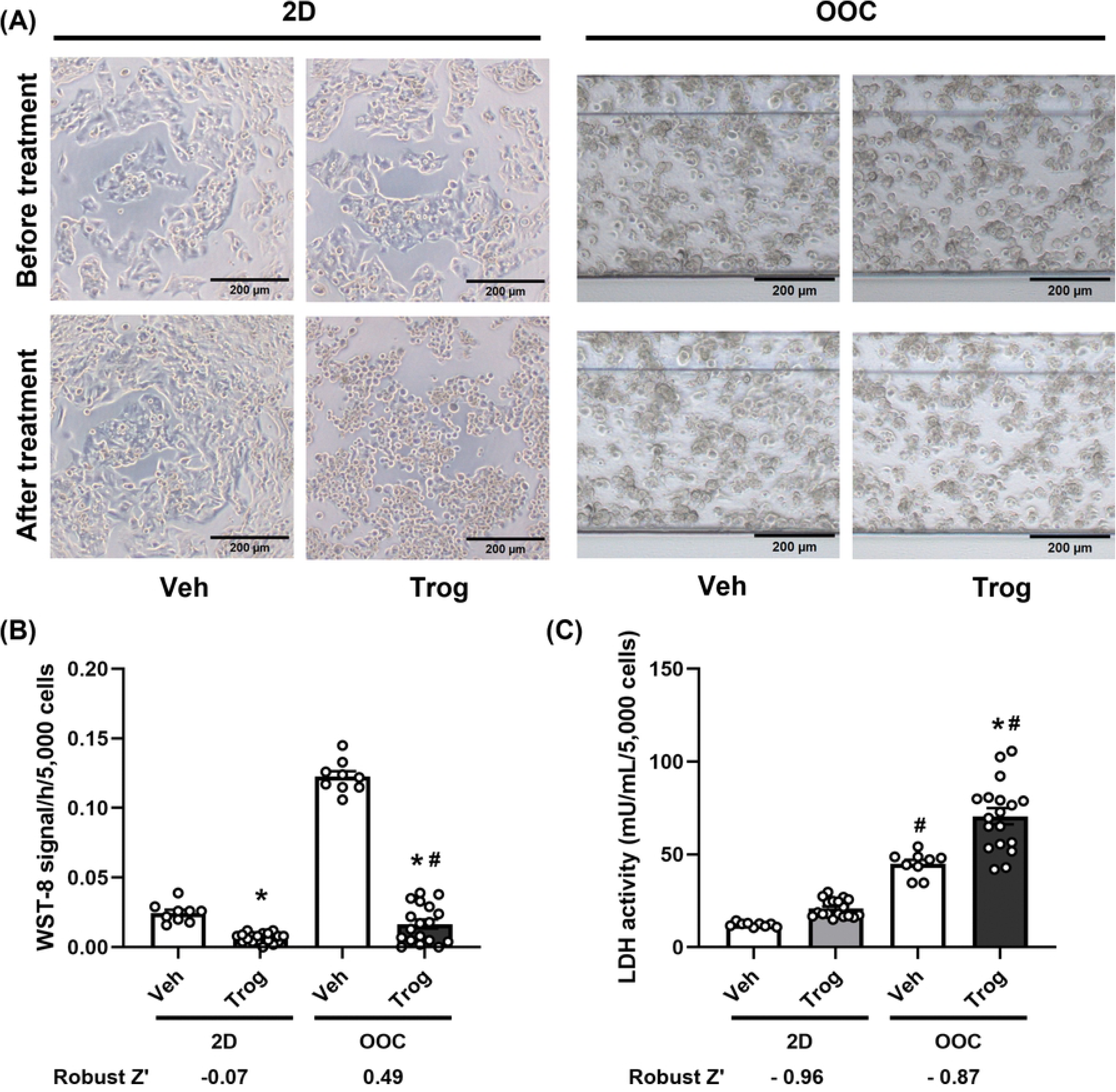
Effects of troglitazone on the viability of HepG2 cells in 2D and OOC cultures. HepG2 cells were treated with 0.1% DMSO (Veh) or 200 µM troglitazone (Trog) for 24 h. (A) Microscopic images of the HepG2 2D and OOC cultures before and after treatment. Scale bar = 200 μm. (B) WST-8 activity per 5,000 seeded cells within an hour of reaction for cell viability. (C) LDH activity per 5,000 seeded cells for cellular damage. The data are expressed as mean ± SEM and analyzed using a one-way ANOVA (n = 9−18) with robust Z’ calculations. * p < 0.05 vs. Veh. ^#^ p < 0.05 vs. 2D Trog.

To evaluate cell viability, we employed the WST-8 assay, which leverages cellular dehydrogenases to convert a tetrazolium substrate into an amber dye with an intensity directly proportional to the cellular metabolic activity [48]. When WST-8 activity was normalized by the seeded cell number, troglitazone treatment significantly reduced the viability of HepG2 cells in the 2D and OOC platforms relative to that of the vehicle-treated cells (Fig 1B). Using the cell culture medium collected after 24-h treatment, cellular damage was assessed by measuring the activity of LDH released into the extracellular environment upon the loss of membrane integrity [59]. Consistent with the WST-8 results, LDH activity was significantly elevated in the conditioned medium of troglitazone-treated HepG2 OOCs compared with that of the vehicle-treated groups (Fig 1C), but not in the samples collected from the 2D cultures (Fig 1C). Based on these concurrent shifts in WST-8 and LDH activities, it appears that the cellular metabolic activities per cell are vastly enhanced in the OOC platform relative to those in the 2D plates. Therefore, these two assays using HepG2 OOCs provide reliable detection of troglitazone-mediated hepatotoxicity; consequently, 200 µM of troglitazone was used as a positive control in subsequent experiments.

To evaluate the reproducibility and robustness of the 2D and OOC cytotoxicity tests for potential high-throughput applications, the Robust Z’-factor was calculated using the dataset from 5−9 independent experiments in which HepG2 OOC cells were treated with vehicle or 200 μM of troglitazone for 24 h. Each experiment consisted of 1−2 technical replicates per treatment (n = 9−18 total replicates). The WST-8 assay dataset from the OOC experiments (but not from the 2D experiments) yielded Robust Z’-factor values > 0.2, indicating acceptable assay quality for high-throughput screening (Fig 1B). Although the LDH assay in both platforms yielded Robust Z’-factor values < 0, indicating an unsuitable baseline assay quality for high-throughput screening (Fig 1C) [54], it remained highly sensitive for cytotoxicity testing, as evidenced by a high statistical power (99.9% for 2D and 99.2% for OOC). These results support the use of the HepG2 OOC system for reliably assessing hepatotoxicity using WST-8 and LDH assays.

### Effects of NATPP0650 (Thiolutin) on the viability and hepatic functions of HepG2 OOCs

The test compounds NATPP0650 and NATPP0437 were previously isolated and characterized by Pansomsuay et al. [16] and Bunbamrung et al. [17], respectively, whereas NATPP0604 was characterized by comparing its ^1^H and ^13^C NMR spectra (S2 Fig) to literature data [47]. The purity of NATPP0650, NATPP0437, and NATPP0604 was confirmed to be > 99% by HPLC analysis (S1 Fig).

To assess the potential hepatotoxicity of NATPP0650, 2D and OOC cultures of HepG2 cells were exposed to vehicle (0.1% DMSO) or various concentrations of NATPP0650 (0.001–50 µM) for 24 h. We found no morphological changes in these cultures following treatment at concentrations ≤ 1 µM, but cells treated at 10 and 50 µM became more rounded and smaller than those in the vehicle-treated groups (Fig 2A and S3 Fig).

**Fig 2.**
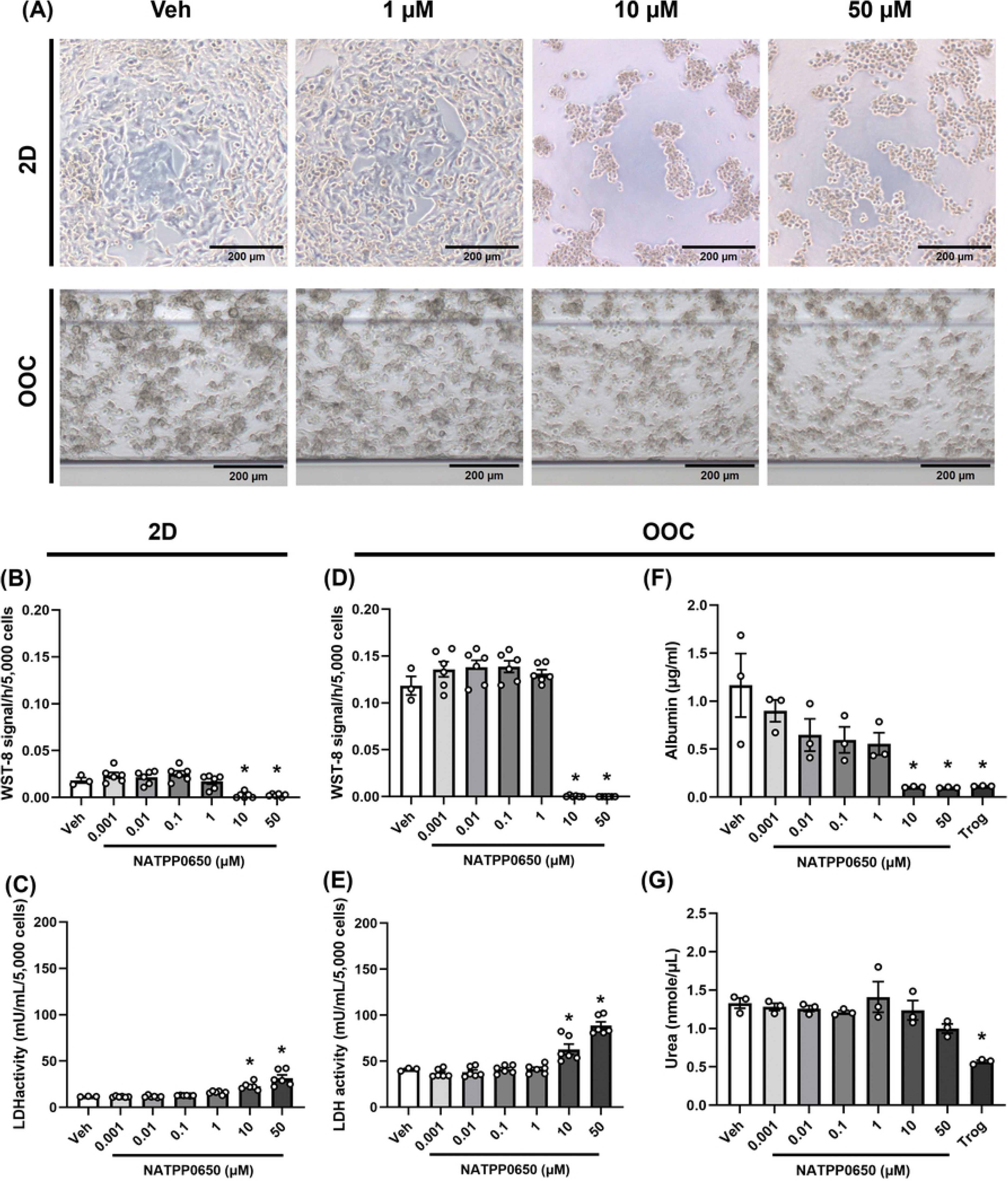
Effects of NATPP0650 (Thiolutin) on the viability and hepatic functions of HepG2 cells cultured in 2D and OOC platforms. HepG2 cells were treated with 0.1% DMSO (Veh), various concentrations of NATPP0650 (0.001−50 µM), or 200 µM troglitazone (Trog) for 24 h. (A) Microscopic images of the HepG2 2D and OOC cultures. Scale bar = 200 μm. (B and D) WST-8 activity for viability in 2D (B) and OOC (D) cultures of HepG2. (C and E) LDH activity for cellular damage in 2D (C) and OOC (E) cultures of HepG2. (F) Albumin production in HepG2 OOCs. (G) Urea production in HepG2 OOCs. The data are expressed as mean ± SEM and analyzed using a one-way ANOVA (n = 3−6). * p < 0.05 vs. Veh.

There were no significant changes in the WST-8 and LDH assay results of HepG2 samples at concentrations ≤ 1 µM (Fig 2B–E). However, 10 and 50 µM of NATPP0650 induced significant declines in WST-8 activity alongside reciprocal increases in LDH leakage relative to the vehicle controls (Fig 2B–E). These results indicate that at concentrations ≤ 1 µM, NATPP0650 affects neither the cell viability nor the cellular integrity of HepG2 cells, regardless of the culture system type.

In the liver, albumin—the most abundant plasma protein—is exclusively synthesized by hepatocytes [60], whereas urea is produced via the urea cycle as the main nitrogenous waste product [61]. These hepatic products are widely used as functional biomarkers in liver OOCs, including HepG2 OOCs [62]. In patients presenting with complications of severe malaria (e.g., malarial hepatitis), the deterioration of hepatic functions is frequently observed in association with decreased serum albumin levels [63, 64]. Hence, an ideal antimalarial drug candidate must exert minimal adverse effects on baseline hepatic functions. To evaluate the potential effects of the candidates on liver functions, we quantified albumin and urea production in the conditioned medium of HepG2 OOCs, which exhibit significantly enhanced albumin and urea production relative to 2D hepatocyte cultures [41, 42]. Treating HepG2 OOCs with NATPP0650 for 24 h reduced albumin production in a concentration-dependent manner (Fig 2F). Notably, albumin levels decreased significantly at doses of 10 and 50 μM, reaching a degree comparable to troglitazone treatment. Conversely, urea production was not significantly altered by NATPP0650 at any tested concentration, whereas troglitazone significantly reduced urea production as expected (Fig 2G). These results suggest that NATPP0650 concentrations ≥ 10 μM exert distinct hepatotoxic effects, as evidenced by alterations in cellular morphology, diminished viability, increased cellular damage, and impaired albumin production without disrupting urea synthesis.

### Effects of NATPP0437 (Cochliodinol) on the viability and hepatic functions of HepG2 OOCs

Next, we examined the toxicological effects of NATPP0437 on HepG2 cells cultured in 2D and OOC platforms. Similar to NATPP0650, we observed morphological changes in the HepG2 cells for both culture platforms after 24-h treatment with NATPP0437 at 10 and 50 µM compared with those in the vehicle-treated controls (Fig 3A and S4 Fig).

**Fig 3.**
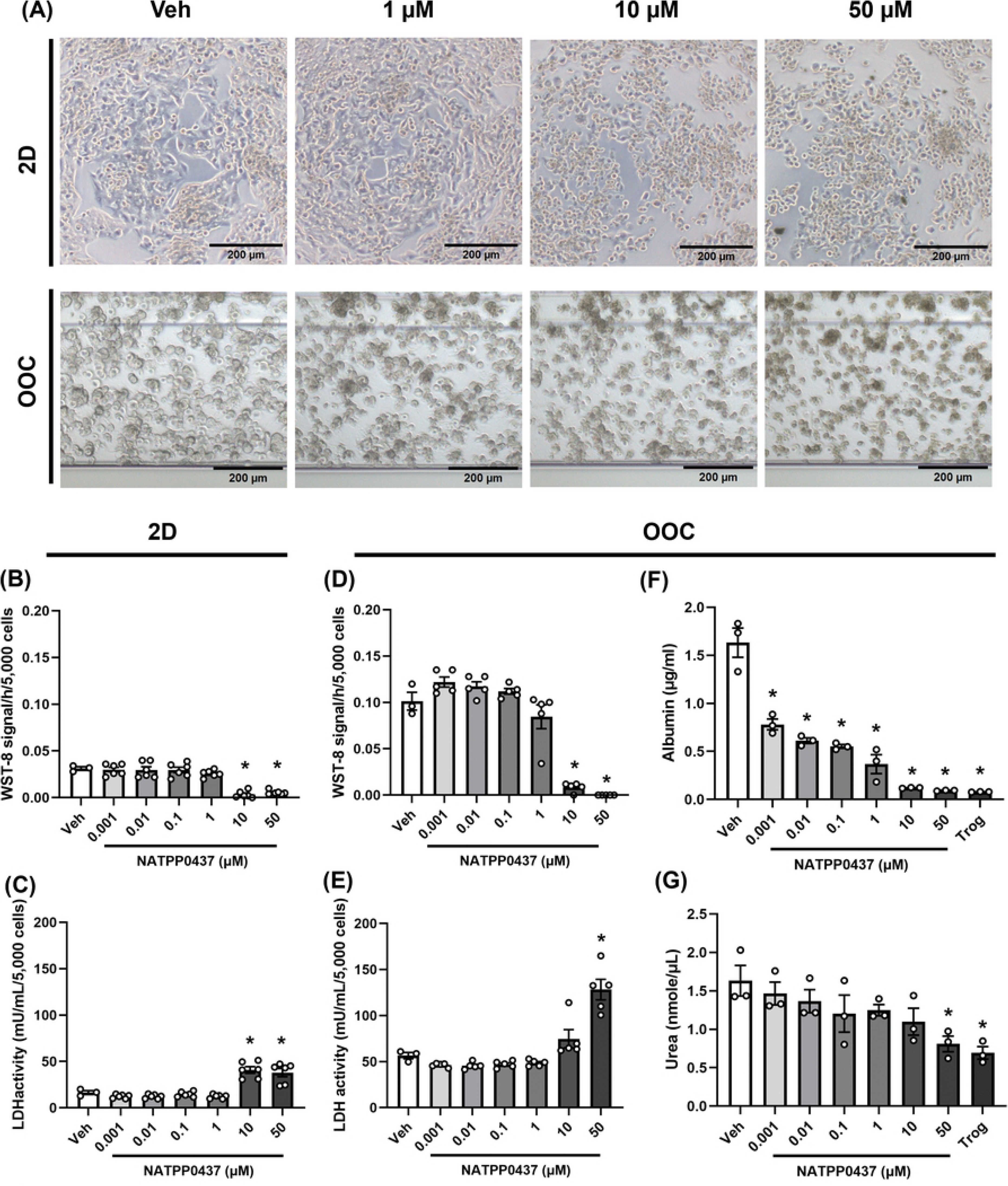
Effects of NATPP0437 (Cochliodinol) on the viability and hepatic functions of HepG2 cells cultured in 2D and OOC platforms. HepG2 cells were treated with 0.1% DMSO (Veh), various concentrations of NATPP0437 (0.001−50 µM), or 200 µM troglitazone (Trog) for 24 h. (A) Microscopic images of the HepG2 2D and OOC cultures. Scale bar = 200 μm. (B and D) WST-8 activity for viability in 2D (B) and OOC (D) cultures of HepG2. (C and E) LDH activity for cellular damage in 2D (C) and OOC (E) cultures of HepG2. (F) Albumin production in HepG2 OOCs. (G) Urea production in HepG2 OOCs. The data are expressed as mean ± SEM and analyzed using a one-way ANOVA (n = 3−6). * p < 0.05 vs. Veh.

These concentrations of NATPP0437 induced significant decreases in WST-8 activity across both systems, compared with the vehicle control (Fig 3B and 3D). Additionally, while LDH activity in the 2D culture medium was significantly increased by 10 and 50 µM of NATPP0437 (Fig 3C), only the highest concentration (50 µM) significantly damaged HepG2 OOCs (Fig 3E). As confirmed by WST-8 and LDH assays in both culture systems, the hepatotoxic nature of NATPP0437 became apparent only at concentrations of 10 and 50 µM.

Functional assessments of the OOCs demonstrated that, similar to NATPP0650, NATPP0437 treatment significantly decreased albumin production in a concentration-dependent manner, with marked reductions occurring at 10 and 50 μM (Fig 3F). Although this concentration-dependent decline was also seen in urea production, this inhibitory effect was significant only when the OOCs were treated with the highest concentration (50 µM) of NATPP0437 relative to the vehicle control (Fig 3G). Collectively, these results suggest that NATPP0437 concentrations ≥10 μM compromise the viability of HepG2 cells in both 2D and OOC formats. Moreover, the albumin-producing capacity of HepG2 OOCs was impaired, whereas urea production remained unaffected except under peak exposure conditions (50 µM).

### Effects of NATPP0604 (4’-hydroxy-MPA) on the viability and hepatic functions of HepG2 OOCs

Unlike the preceding two compounds, NATPP0604 did not alter cellular morphology at any tested concentration (Fig 4A and S5 Fig).

**Fig 4.**
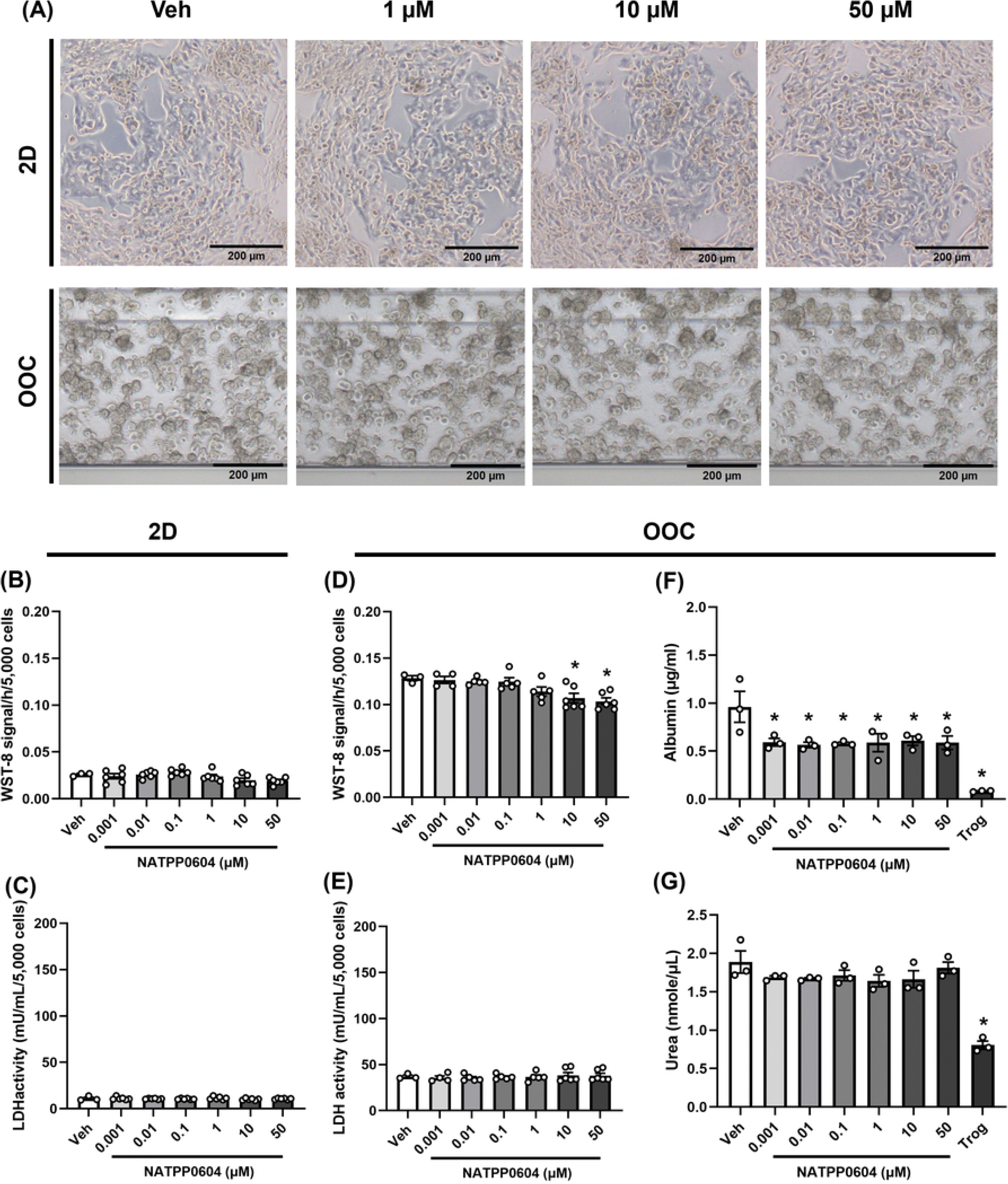
Effects of NATPP0604 (4’-hydroxy-MPA) on the viability and hepatic functions of HepG2 cells cultured in 2D and OOC platforms. HepG2 cells were treated with 0.1% DMSO (Veh), various concentrations of NATPP0604 (0.001−50 µM), or 200 µM troglitazone (Trog) for 24 h. (A) Microscopic images of the HepG2 2D and OOC cultures. Scale bar = 200 μm. (B and D) WST-8 activity for viability in 2D (B) and OOC (D) cultures of HepG2. (C and E) LDH activity for cellular damage in 2D (C) and OOC (E) cultures of HepG2. (F) Albumin production in HepG2 OOCs. (G) Urea production in HepG2 OOCs. The data are expressed as mean ± SEM and analyzed using a one-way ANOVA (n = 3−6). * p < 0.05 vs. Veh.

This result was supported by both the WST-8 (Fig 4B and 4D) and LDH assays (Fig 4C and 4E), which showed that metabolic activities and membrane integrity remained mostly unaffected in the presence of NATPP0604 across both culture platforms. Consistently, urea production was unaffected by NATPP0604 at all concentrations (Fig 4G). In contrast, we found steady, concentration-independent decreases in albumin production for HepG2 OOCs exposed to NATPP0604 relative to those treated with vehicle (Fig 4F). However, the extent of this NATPP0604-induced albumin reduction was less pronounced than that induced by NATPP0650 (Fig 2F) and NATPP0437 (Fig 3F), particularly at 10 and 50 µM. These results suggest that NATPP0604 exhibits marginal acute hepatotoxicity, along with limited detrimental impact on albumin production, compared with NATPP0650 and NATPP0437.

To extend our understanding of the toxicity profiles for the three compounds within the vasculature—where pathological malaria–endothelial interactions induce barrier integrity disruption, inflammatory responses, vascular leakage, and cerebral edema [65–67]—we employed a 2-lane OrganoPlate lined with human umbilical vein endothelial cells (HUVECs). This cell type has been shown to closely reproduce key features of vascular endothelium in vitro [65, 66]. HUVEC-based OOC models have been developed by other groups [68] and previously validated by our group [69–71]. After treating HUVEC OOCs with either 0.1% DMSO (vehicle) or various concentrations (0.05, 0.5, 5, and 50 μM) of NATPP0650, NATPP0437, or NATPP0604 for 24 h, we evaluated their acute toxicity via bright-field imaging and LDH quantification. NATPP0650 at concentrations ≥ 0.5 μM caused morphological changes and cellular detachment from the collagen ECM (S6 Fig). This damage was consistent with significant increases in LDH activity within the conditioned medium collected from HUVEC OOCs exposed to the corresponding concentrations of NATPP0650 (S6 Fig). While less toxic than NATPP0650, 5 and 50 μM of NATPP0437 also significantly increased LDH activity when the morphological abnormality was observed in HUVEC OOCs (S7A and S7B Figs). Conversely, NATPP0604 induced no significant cytotoxicity based on structural morphology or LDH leakage relative to the vehicle control in HUVEC OOCs (S8A and S8B Figs). Our findings indicate that the cytotoxicity profiles of the tested compounds vary between the hepatic and endothelial cell types. In HUVECs, 0.5 μM of NATPP0650 or 5 μM of NATPP0437 was sufficient to provoke significant LDH release, whereas HepG2 cells remained insensitive to these treatments at concentrations < 10 μM. Notably, NATPP0604 displayed no detectable cytotoxicity in either OOC model within the tested concentration ranges, suggesting its superior safety profile over the other two compounds.

### Effects of prolonged treatment with NATPP0650 (Thiolutin) on the viability and albumin production of HepG2 OOCs

Treating malaria patients with ACTs, the current gold standard, for a duration of 3–7 days is a standard procedure for achieving complete parasitic remission [1, 31–34]. To discover novel antimalarial drug candidates that also fulfill this operational window, conventional 2D culture systems may be insufficient owing to their inability to maintain long-term cultivation [37]. In contrast, HepG2 OOCs can sustain viability for at least 14 days and have been implemented in toxicological evaluations of acetaminophen, palmitate, and plant extracts [41–43], making them suitable for studying the effects of prolonged exposure. To this end, we treated HepG2 OOCs with medium containing 0.1% DMSO (vehicle), various concentrations of NATPP0650 (0.001–50 µM), or 200 μM of troglitazone, with daily media replacement for 7 days. We observed evident cell shrinkage and the loss of cluster formation after 7 days of repeated treatment with 10 or 50 µM NATPP0650 (Fig 5A and S9 Fig).

**Fig 5.**
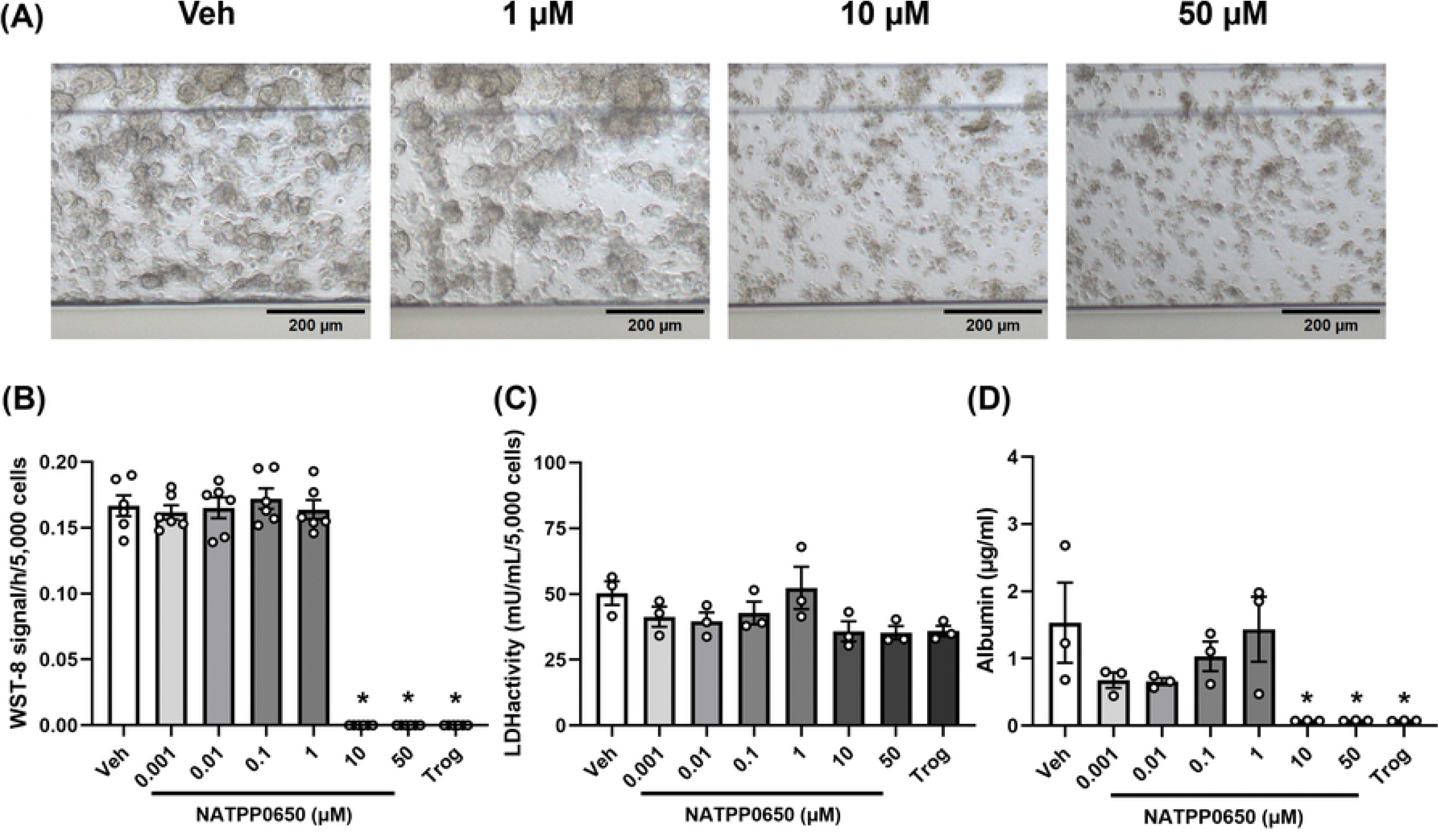
Effects of prolonged treatment with NATPP0650 (Thiolutin) on the viability and albumin production of HepG2 OOCs. HepG2 cells were treated with 0.1% DMSO (Veh), various concentrations (0.001−50 µM) of NATPP0650, or 200 µM troglitazone (Trog) repeatedly for 7 days. (A) Microscopic images of the HepG2 OOCs after 7-day repeated treatment. Scale bar = 200 μm. (B) WST-8 activity for viability. (C) LDH activity for cellular damage. (D) Albumin production. The data are expressed as mean ± SEM and analyzed using a one-way ANOVA (n = 3−6). * p < 0.05 vs. Veh.

Correspondingly, WST-8 assay results showed significant and robust decreases in HepG2 OOC viability following the 7-day exposure to NATPP0650 at 10 and 50 µM but not at 0–1 µM (Fig 5B). Conversely, negligible changes in extracellular LDH activity were detected in the conditioned medium of NATPP0650-treated HepG2 OOCs across all concentrations (Fig 5C). This presumably occurs because the severe cell death happened rapidly within the first 24–48 hours of the long-term regimen, leaving no viable, intact cells left to leak fresh LDH by day 7. As observed in the acute 24-h treatment (Fig 2F), albumin production was completely suppressed by the 7-day repeated treatment with NATPP0650 at 10 or 50 μM (Fig 5D).

### Effects of prolonged treatment with NATPP0437 (Cochliodinol) on the viability and albumin production of HepG2 OOCs

Consistent with the long-term toxicity profiles observed for NATPP0650 (Fig 5), treating HepG2 OOCs with NATPP0437 (10 and 50 µM) for 7 days caused distinct morphological changes (Fig 6A and S10 Fig) and significant reductions in WST-8 activity (Fig 6B) and albumin production (Fig 6D), without affecting LDH activity (Fig 6C).

**Fig 6.**
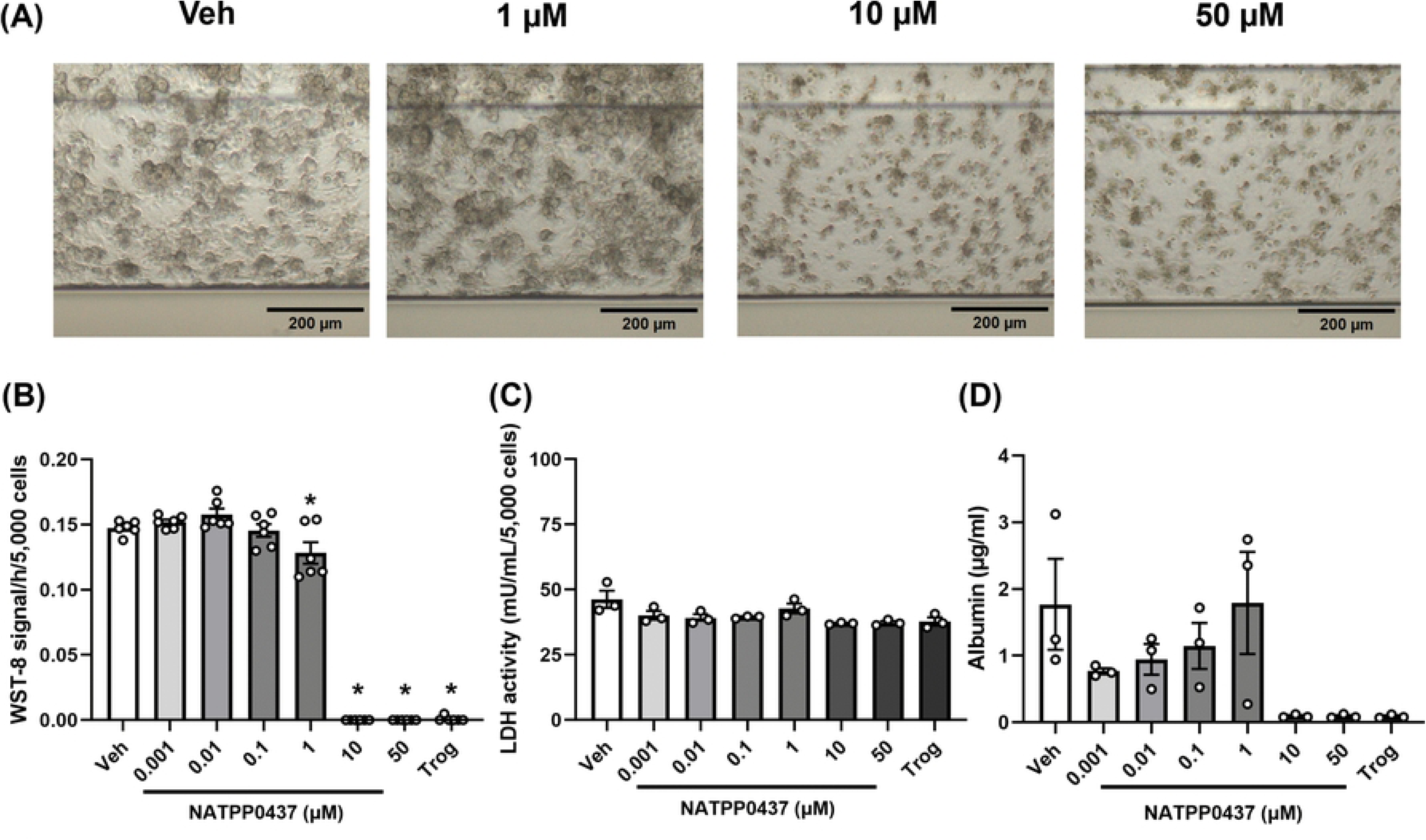
Effects of prolonged treatment with NATPP0437 (Cochliodinol) on the viability and albumin production of HepG2 OOCs. HepG2 cells were treated with 0.1%DMSO (Veh), various concentrations (0.001−50 µM) of NATPP0437, or 200 µM troglitazone (Trog) repeatedly for 7 days. (A) Microscopic images of the HepG2 OOCs after 7-day repeated treatment. Scale bar = 200 μm. (B) WST-8 activity for viability. (C) LDH activity for cellular damage. (D) Albumin production. The data are expressed as mean ± SEM and analyzed using a one-way ANOVA (n = 3−6) * p < 0.05 vs. Veh.

These results indicate that treatment with NATPP0437 at concentrations > 10 µM compromise both the viability and function of HepG2 OOCs during long-term culture, analogous to the toxicity characteristics of NATPP0650. In summary, both NATPP0650 and NATPP0437 significantly impair the structural integrity and metabolic functionality of HepG2 OOCs following a 7-day repeated-exposure regimen at concentrations > 10 µM.

### Effects of prolonged treatment with NATPP0604 (4’-hydroxy-MPA) on the viability and albumin production of HepG2 OOCs

Finally, we examined the potential long-term toxicity of NATPP0604 in HepG2 OOCs. Unlike NATPP0650 and NATPP0437, no morphological changes were found in HepG2 OOCs following the 7-day treatment with 10 or 50 µM of NATPP0604 compared with those following vehicle treatment (Fig 7A and S11 Fig).

**Fig 7.**
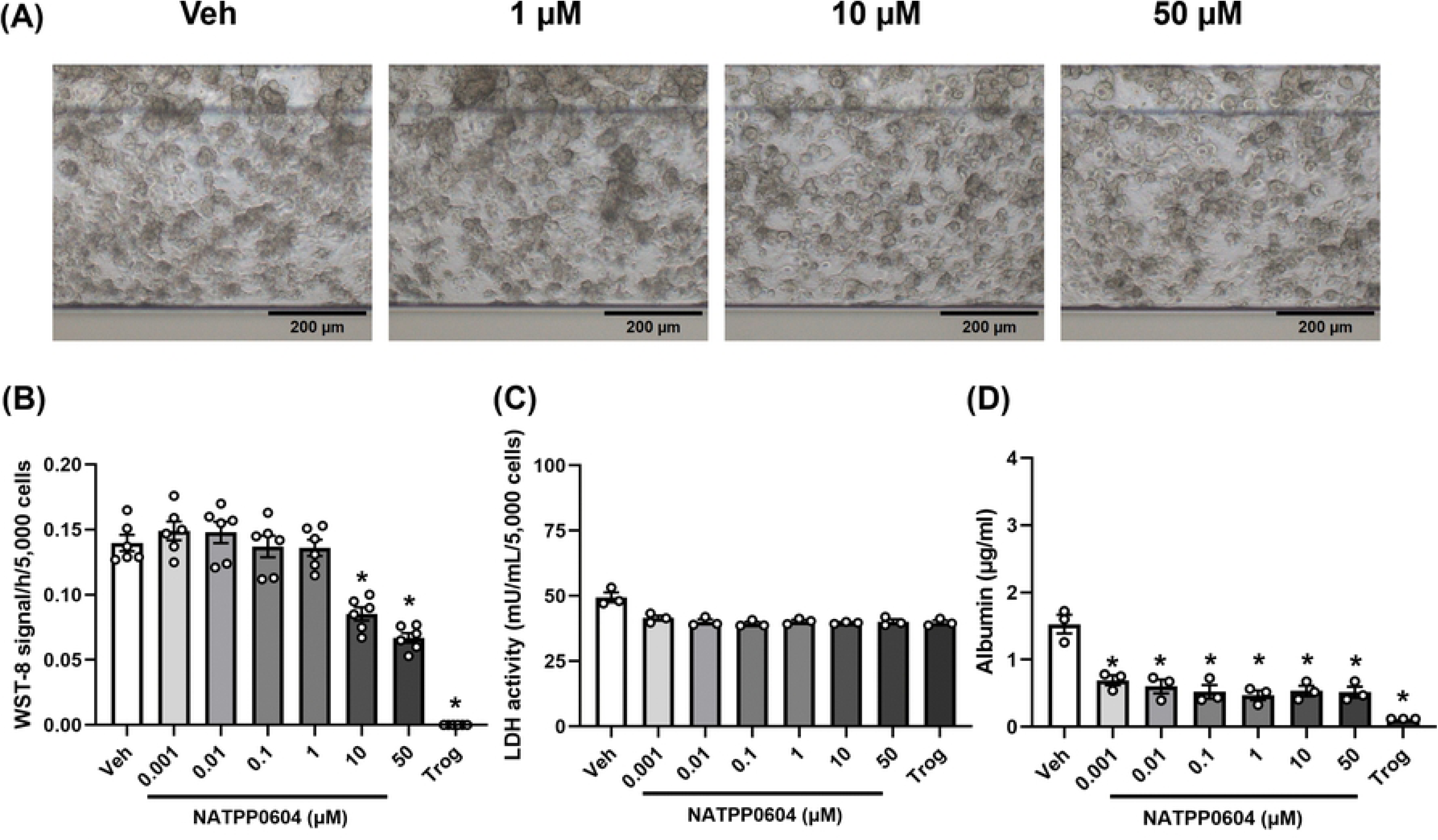
Effects of prolonged treatment with NATPP0604 (4’-hydroxy-MPA) on the viability and albumin production of HepG2 OOCs. HepG2 cells were treated with 0.1%DMSO (Veh), various concentrations (0.001−50 µM) of NATPP0604, or 200 µM troglitazone (Trog) repeatedly for 7 days. (A) Microscopic images of the HepG2 OOCs after 7-day repeated treatment. Scale bar = 200 μm. (B) WST-8 activity for viability. (C) LDH activity for cellular damage. (D) Albumin production. The data are expressed as mean ± SEM and analyzed using a one-way ANOVA (n = 3−6) * p < 0.05 vs. Veh.

Additionally, while NATPP0650 and NATPP0437 both completely eliminated OOC viability at these upper thresholds (Figs 5B and 6B), 10 and 50 µM of NATPP0604 decreased cell viability by only 39% and 52%, respectively relative to the vehicle controls (Fig 7B). Consistent with previous observations under daily media replacement, no significant changes in LDH activity were observed (Fig 7C). Meanwhile, NATPP0604 induced a uniform, significant reduction in albumin production across all tested concentrations (Fig 7D), unlike the dose-dependent inhibitory profiles observed for NATPP0650 and NATPP0437 (Figs 5D and 6D). These results suggest that NATPP0604 possesses lower hepatotoxicity than the other two compounds, even during repeated exposure to high concentrations (10 and 50 µM) for 7 days.

Collectively, comparing the 24-h and 7-day toxicological timelines of the three compounds using HepG2 OOCs provides key insights into candidate safety. The 7-day repeated treatment with NATPP0650 or NATPP0437 at 10 and 50 µM decreased the viability and albumin production of the HepG2 OOCs as severely as their 24-h treatments. However, the cytotoxicity of NATP0604 at 10 and 50 µM was revealed only by the 7-day procedure, remaining undetected in the 24-h experiments. These outcomes support the use of the HepG2 OOCs for the long-term toxicity screening of bioactive compounds in vitro.

## Discussion

In this study, we assessed the hepatotoxic potential of three drug candidates developed for malaria treatment using the HepG2 OOC system. For comparison, conventional 2D cultures of HepG2 cells were used in the 24-h toxicity studies to evaluate morphological changes, metabolic viability, and cellular damage. Our findings revealed several key toxicological insights in the acute screening phase: 1) NATPP0650 and NATPP0437 exhibited hepatotoxicity at high concentrations (10 and 50 µM), as indicated by cellular shrinkage, reduced viability, membrane leakage, and compromised functional synthesis; 2) NATPP0604 exhibited no hepatotoxicity at any concentration, showing only a mild, concentration-independent reduction in albumin production; 3) no apparent variations occurred in the viability or cellular damage assessment between 2D and OOC cultures of HepG2 cells during 24-h exposure; and 4) in HUVEC OOCs, NATPP0604 showed no acute cytotoxicity, whereas NATPP0437 (5 and 50 µM) and NATPP0650 (≥ 0.5 µM) exhibited significant toxicity. In the long-term (7-day) OOC studies, we found that 5) NATPP0650 and NATPP0437 remained non-toxic up to 1 µM but displayed cumulative toxicity at 10 and 50 µM, whereas 6) NATPP0604 exhibited latent toxicity at 10 and 50 µM.

While conventional in vitro 2D liver models remain useful for initial drug safety evaluation [30], their utility is constrained by the absence of a supporting ECM scaffold, which compromises cellular stability and culture longevity [30]. Furthermore, without fluidic flow, hepatic functions are not fully maintained over time [72–74]. In the present study, the reliability of assays (e.g., WST-8) in 2D HepG2 monolayers was undermined by unintended cell detachment during the repetitive washing steps required to remove residual chemicals (S12 Fig). This is a well-known vulnerability of adherent 2D monolayers, which are highly susceptible to mechanical disruption and changes in microenvironmental conditions during chemical exposure [75]. These limitations are effectively mitigated in OOC platforms [30, 76, 77]. By providing an in vivo-like microenvironment, OOCs recapitulate physiological cell–cell and cell–ECM interactions [78, 79], while dynamic microfluidic perfusion facilitates the continuous delivery of nutrients and removal of metabolic waste [78, 79]. Moreover, the embedded ECM shields cells from direct fluid shear stress (e.g., medium flow and washing), thereby ensuring more consistent cell densities and reliable metabolic readouts [80].

We initially benchmarked the sensitivity of our HepG2 OOC system using troglitazone, a clinical therapeutic known to induce mitochondrial dysfunction and oxidative stress in humans [58]. The results indicated that troglitazone significantly reduced cell membrane integrity in the OOC model but not in the 2D culture. This increased sensitivity to troglitazone in the OOC system may originate from the biomimetic microenvironment, which supports enhanced drug uptake and amplifies the response to troglitazone [81, 82]. Additionally, the OOC platform exhibited heightened baseline metabolic activities, as reflected in the basal and troglitazone-treated levels of the WST-8 and LDH readouts. Collectively, these results reaffirm that fluid flow and a three-dimensional structure can enhance the drug sensitivity and metabolic activity of liver OOCs [30, 36, 37, 76, 82].

With the global emergence of ART resistance [1, 7–9], searching for structurally novel antimalarial scaffolds remains a high priority. *Streptomyces* species have been shown to produce various metabolites with parasiticidal properties against selected *Plasmodium* strains [83]. NATPP0650 isolated from *Streptomyces marinisediminis* JHD1^T^ exhibited an IC_50_ value of 5.92 µM against the multidrug-resistant *P. falciparum* K1 strain [16], falling within the established threshold for good antiplasmodial activity (IC_50_ of 1−20 µM) [18]. Similar to the genus *Streptomyces*, the genus *Chaetomium* is known to produce metabolites with antimalarial bioactivities [84, 85]. However, azaphilones derived from *C. globosum* can exert harmful effects on HepG2 cells [86]. In accordance with these reports, NATPP0437—an azaphilone produced from *C. globosum*—demonstrated antimalarial activity (IC_50_ of 4.39 µM against *P. falciparum* K1 strain) [17] and cytotoxic properties when evaluated in HepG2 and HUVEC OOC systems at moderate-to-high concentrations. The most promising candidate in the present study, NATPP0604 (IC_50_ of 2.11 µM), was isolated from *Penicillium parvum* BCC 75476. Future studies are needed to further characterize this fungal strain, considering that it may produce other antimalarial and bioactive substances.

Historically, the most notable sources of antimalarials have been plants, such as *Artemisia annua* and *Cinchona*, from which artemisinin and quinine were originally isolated, respectively [87]. The synthetic derivatives of artemisinin, such as artemether and artesunate, have been widely used as components of ACTs [87, 88] and remain the gold standard for treating uncomplicated falciparum malaria [31]. This clinical dominance is supported by IC_50_ values of 1–5 nM for various ART compounds against the *P. falciparum* K1 strain during the asexual blood stage [89, 90]. In a murine malaria model, several ACT regimens reduce parasite burden while mitigating parasite-induced hepatorenal toxicity [91]. Similarly, an observational cohort study in Ethiopia reported that malaria infection correlates with elevated liver enzymes and anomalous lipid profiles, which were both normalized after treatment with antimalarial drugs [90]. In contrast, while the association of ACTs with transient ALT/AST elevations has also been reported, these anomalies typically resolve post-recovery without progressing to clinical liver injury [92]. These lines of evidence indicate that ACTs can provide rapid parasite clearance with a favorable hepatic safety profile. Consequently, novel antimalarial agents should achieve parasiticidal plasma concentrations comparable to those of existing ACTs while demonstrating equivalent or lower rates of hepatotoxicity.

The SI has been commonly used to demonstrate the potential of compounds that effectively inhibit parasite growth (i.e., low IC_50_ values) without inducing host cytotoxicity (i.e., high CC_50_ values), including hepatotoxicity [52]. Based on standard SI calculations (SI = CC_50_/IC_50_), drug candidates with an SI value > 10 should offer a more potent antimalarial and safer therapy [51–53]. For instance, the SI of the standard antimalarial artesunate was 42,475 (calculated as the ratio of the CC_50_ in Vero cells to the IC_50_ against the chloroquine-sensitive *P. falciparum* 3D7 strain), indicating highly specific antiparasitic action [93]. When evaluated against HepG2 hepatocytes, artesunate maintained a robust, albeit lower, SI of 7,000 [52]. This variation in SI values highlights the critical importance of cell-type selection when establishing therapeutic safety margins. Furthermore, the use of HepG2 hepatocytes is highly significant for the clinically predictive safety evaluation of antimalarial compounds [51], given that the liver serves as the primary site for initial parasite replication. Although all three candidates evaluated in the present study exhibited low-micromolar IC_50_ values, their SI profiles based on the 24-h OOC results differed markedly. The calculated SI values for NATPP0650 and NATPP437 were 0.51 and 0.50, respectively, indicating that these compounds provoke human liver cell toxicity at concentrations below their effective antiparasitic thresholds [53]. This suggests that their antimalarial effects may be a secondary consequence of general cellular stress or “promiscuous” toxicity, which would likely disqualify them from further development [51, 53]. Conversely, NATPP0604 exhibited CC_50_ values > 50 µM, yielding a high SI > 23.7, superior to the other two compounds investigated herein. Although the absolute efficacy and SI of these three microbe-derived compounds are lower than those of some plant-derived compounds [94], this study corroborated the short-term safety of NATPP0604 in HepG2 and HUVEC OOCs, as well as the long-term safety of NATPP0604 in HepG2 OOCs. These insights encourage further development of NATPP0604 through targeted chemical modification to enhance its efficacy [95–97].

It is common to find drug candidates that show little or no cytotoxicity after 24 h of exposure yet display toxicity during long-term exposure [98]. The treatment duration of antimalarial candidates in preclinical settings should correspond to established antimalarial regimens; standard ACTs require administration for at least 3 days [1, 31–34]. Consequently, traditional 2D models subjected to 24-h exposure fail to fully simulate the clinical reality. In vitro hepatotoxicity has been reported in HepG2-derivative cells repeatedly exposed to several antimalarial agents for up to 72 h [12]. To bridge this translational gap, the present study evaluated the long-term effects of three compounds using HepG2 OOCs capable of sustaining 7-day repeated-exposure toxicity assessments [43]. We found that although long-term exposure to NATPP0604 can harm liver cells, concentrations < 10 µM—at which NATPP0604 exhibits antimalarial effects (IC_50_ = 2.11 µM)—remain safe over 7 days. However, because albumin production was partially suppressed even at the lowest tested concentration, plasma albumin levels should be carefully monitored in patients during future clinical trials involving this compound.

We also observed the uncoupling of the two hepatic biomarkers: while albumin synthesis declined sharply upon exposure, urea production remained largely unaffected, even at high concentrations sufficient to significantly deteriorate cell viability. This phenomenon can be attributed to albumin’s role as a negative acute-phase protein, whose expression decreases rapidly when liver cells encounter chemical toxins [99, 100]. In collagen sandwich cultures, exposing rat hepatocytes to classic hepatotoxic compounds, such as aflatoxin and methyl methane sulfonate, caused a more significant and rapid decline in albumin production than in urea synthesis [100]. Aligned with this precedent, our findings demonstrate that albumin is a more sensitive functional biomarker for drug-induced liver injury than urea because it exhibits a greater decline following exposure to the test compounds. Accordingly, our data also highlight the utility of HepG2 OOCs for exploring the underlying mechanisms driving the uncoupling of albumin and urea pathways.

*P. falciparum* is known to compromise endothelial barrier integrity through direct cytoadhesion and immune activation [67], contributing to dysfunction in the blood–brain barrier [101–103], pulmonary microvasculature [104–106], and renal capillaries [103, 107]. Notably, standard artemisinin-based therapies can negatively impact endothelial proliferation or fail to protect barrier integrity. For instance, dihydroartemisinin has been shown to inhibit microvascular endothelial cell growth at therapeutic concentrations [108], whereas artemisinin fails to prevent further barrier damage in cerebral malaria models [109]. Hence, it is critical to ensure that novel antimalarial candidates do not exacerbate this pre-existing endothelial vulnerability. Similar to dihydroartemisinin’s detrimental action on endothelial cells, NATPP0650 and NATPP0437 at moderate-to-high concentrations showed significant toxicity in HUVEC OOCs within 24 h. Conversely, no toxicity was present in HUVEC OOCs after 24-h exposure to NATPP0604. Collectively, NATPP0604 demonstrated the least toxicity in both HepG2 and HUVEC OOCs among the three candidates, offering the safety assurance for further drug development. Nevertheless, the long-term toxicity assessment in HUVEC OOCs remains to be explored. While integrated malaria-OOC platforms incorporating hepatocytes, HUVECs, splenocytes, and red blood cells have been developed to capture the systemic nature of *P. falciparum* infection and off-target toxicity of antimalarial compounds [110], their complexity often complicates the interpretation of cellular-level interactions and can introduce issues with reproducibility [111]. To balance these trade-offs, our independent assessments in HepG2 and HUVEC OOC models offer a focused, high-resolution evaluation of cell type-specific toxicity.

We acknowledge some limitations specific to HepG2 OOCs. First, our current system relies on a monoculture and does not account for the role of non-parenchymal immune cells in mediating toxic responses [112]. Future integration of co-culture configurations in an OOC platform may facilitate mechanistic studies of immune-mediated, drug-induced liver injury [112]. Second, our system uses gravity-driven fluid flow that reverses direction every 8 min, resulting in a bidirectional flow profile. *In vivo*, hepatic blood flow is strictly unidirectional, which ensures optimal waste removal and a continuous supply of fresh nutrients [113]. Although it does not perfectly replicate the *in vivo* setting, the bidirectional flow present in our OOC model provides a superior physiological microenvironment compared with traditional 2D cultures. This is evidenced by significantly higher metabolic activity via WST-8-related pathways and enhanced LDH release when normalized to an equivalent seeded cell number of 5,000 cells. These metabolic alterations may be attributed to the improved cell–cell and cell–ECM interactions under fluidic flow, which has frequently been observed in microphysiological systems [30, 36, 37, 76, 80, 82]. Third, to the best of our knowledge, there are no reports on the specific metabolic pathways and potential cytochrome P450 (CYP)-mediated biotransformation of these three compounds. It has been shown that HepG2 cells have limited basal expression and inducibility of key CYP enzymes relative to those of primary human hepatocytes [114, 115], despite enhanced CYP metabolism in their 3D cultures [41, 116–118]. Therefore, it remains unclear whether the observed hepatotoxicity in the present study is predominantly owing to the parent compounds or OOC-generated metabolites. Future comprehensive studies using mass spectrometry or metabolic profiling are warranted to characterize the metabolic fate of these three compounds. Finally, we assessed the potential toxicity of the three compounds using HepG2 cells that had not been infected with *Plasmodium* parasites. This experimental design does not exclude the possibility that infected HepG2 cells may be more prone to exhibiting toxic effects than non-infected ones. Further studies, particularly those optimizing NATPP0604, should use *Plasmodium* parasite-infected liver OOCs to better simulate a clinically relevant pathophysiological environment.

## Conclusions

This study showed that NATP0604 possesses the most favorable hepatic and endothelial safety profiles among the tested microbial compounds, characterized by an excellent SI value (> 23.7), which represents a more potent antimalarial and a safer therapeutic window [51–53]. Consequently, NATPP0604 may serve as a highly valuable chemical scaffold for hit-to-lead structural optimization, where future semi-synthetic modifications can aim to aggressively drive down the anti-plasmodial IC_50_ into the nanomolar range, thereby exponentially expanding its SI to match modern ACT standards. Unlike conventional 2D cultures, HepG2 OOCs can maintain cellular functionality during prolonged cultivation, enabling the sensitive detection of delayed toxic effects following repeated exposure. Because NATP0604 exhibited robust long-term safety—despite manageable disturbances in baseline albumin production—the findings support its potential for further development as an antimalarial drug.

## Acknowledgments

We thank Miss Anunyaporn Phungsom and Miss Mintra Kwathai for providing the supplementary data of the toxicity assessment using HUVEC OOCs.

## Supporting information captions

**S1 Fig. The high-performance liquid chromatography (HPLC) results of (A) NATPP0650, (B) NATPP0437, and (C) NATPP0604.**

**S2 Fig. NMR spectroscopy of the NATPP0604 compound.** (A) ^1^H NMR spectrum of NATPP0604 in DMSO-*d_6_* (400 MHz). (B) ^13^C NMR spectrum of NATPP0604 in DMSO-*d_6_* (100 MHz).

**S3 Fig. Effect of NATPP0650 (Thiolutin) on the viability of 2D and OOC culture of HepG2 cells.** HepG2 cells were treated with 0.1%DMSO (Veh), various concentrations (0.001−0.1 µM) of NATPP0650 for 24 h. Microscopic images captured at 5 × magnification of the HepG2 cells both 2D and OOC platforms after 24 h treatment. Scale bar = 200 µm.

**S4 Fig. Effect of NATPP0437 (Cochliodinol) on the viability of 2D and OOC culture of HepG2 cells.** HepG2 cells were treated with 0.1%DMSO (Veh), various concentrations (0.001−0.1 µM) of NATPP0437 for 24 h. Microscopic images captured at 5 × magnification of the HepG2 cells both 2D and OOC platforms after 24 h treatment. Scale bar = 200 µm.

**S5 Fig. Effect of NATPP0604 (4’-hydroxy-MPA) on the viability of 2D and OOC culture of HepG2 cells.** HepG2 cells were treated with 0.1%DMSO (Veh), various concentrations (0.001−0.1 µM) of NATPP0604 for 24 h. Microscopic images captured at 5 × magnification of the HepG2 cells both 2D and OOC platforms after 24 h treatment. Scale bar = 200 µm.

**S6 Fig. Effect of NATPP0650 (Thiolutin) on the viability of 3-D HUVECs.** HUVEC cells were treated with 0.1%DMSO (Veh), various concentrations (0.05−50 µM) of NATPP0650 for 24 h. (A) Microscopic images at 5 × magnification of the 3-D HUVECs after 24 h treatment. (B) LDH activity for cellular damage in 3-D HUVECs. The data are expressed as mean ± SEM and analyzed using a one-way ANOVA (n = 3−6). * *p* < 0.05 vs. Veh.

**S7 Fig. Effect of NATPP0437 (Cochliodinol) on the viability of 3-D HUVECs.** HUVEC cells were treated with 0.1%DMSO (Veh), various concentrations (0.05−50 µM) of NATPP0437 for 24 h. (A) Microscopic images at 5 × magnification of the 3-D HUVECs after 24 h treatment. (B) LDH activity for cellular damage in 3-D HUVECs. The data are expressed as mean ± SEM and analyzed using a one-way ANOVA (n = 3−6). * *p* < 0.05 vs. Veh.

**S8 Fig. Effect of NATPP0604 (4’-hydroxy-MPA) compound on the viability of 3-D HUVECs.** HUVEC cells were treated with 0.1%DMSO (Veh), various concentrations (0.05−50 µM) of NATPP0604 for 24 h. (A) Microscopic images at 5 × magnification of the 3-D HUVECs after 24 h treatment. (B) LDH activity for cellular damage in 3-D HUVECs. The data are expressed as mean ± SEM and analyzed using a one-way ANOVA (n = 3−6). * *p* < 0.05 vs. Veh.

**S9 Fig. Effect of NATPP0650 (Thiolutin) on the viability of HepG2 OOCs.** HepG2 cells were treated with 0.1%DMSO (Veh), various concentrations (0.001−0.1 µM) of NATPP0650 repeatedly for 7 Days. Microscopic images captured at 5 × magnification of HepG2 OOCs after 7 days of treatment. Scale bar = 200 µm.

**S10 Fig. Effect of NATPP0437 (Cochliodinol) on the viability of HepG2 OOCs.** HepG2 cells were treated with 0.1%DMSO (Veh), various concentrations (0.001−0.1 µM) of NATPP0437 repeatedly for 7 Days. Microscopic images captured at 5 × magnification of HepG2 OOCs after 7 days of treatment. Scale bar = 200 µm.

**S11 Fig. Effect of NATPP0604 (4’-hydroxy-MPA) on the viability of HepG2 OOCs.** HepG2 cells were treated with 0.1%DMSO (Veh), various concentrations (0.001−0.1 µM) of NATPP0604 repeatedly for 7 Days. Microscopic images captured at 5 × magnification of HepG2 OOCs after 7 days of treatment. Scale bar = 200 µm.

**S12 Fig. Comparison of 2D HepG2 before and after washing processes to remove the residual chemicals.** Cells that were exposed to Veh (0.1%DMSO) showed almost identical before and after the washing processes. Cells that were exposed to Trog (troglitazone) showed a notable loss of cells after washing processes.

